# ASC specks and α-synuclein amplify microglial NLRP3 inflammasome activation and co-localize with Lewy body pathology

**DOI:** 10.64898/2026.10.02.756339

**Authors:** Karan Thakur, Christine Péladeau, Melissa Hewitt, Klaudia Baumann, David Cai, John M. Woulfe, Julianna J. Tomlinson, Michael G. Schlossmacher, Jagdeep K. Sandhu

**Affiliations:** Department of Cellular and Molecular Medicine, University of Ottawa, Ottawa, ON, Canada; Human Health Therapeutics Research Centre, National Research Council Canada, Ottawa, ON, Canada; Neuroscience Program, Ottawa Hospital Research Institute, Ottawa, ON, Canada; Department of Pathology and Laboratory Medicine, The Ottawa Hospital, Ottawa, ON Canada; Division of Neurology, Department of Medicine, The Ottawa Hospital, Ottawa, ON, Canada; Centre for Infection, Immunity and Inflammation (CI3), Faculty of Medicine, University of Ottawa, Ottawa, ON, Canada; Department of Biochemistry, Microbiology, and Immunology, University of Ottawa, Ottawa ON, Canada

**Author notes:** Correspondence: Jagdeep K. Sandhu (;) and Michael G. Schlossmacher.

**Keywords:** synucleinopathies, Parkinson’s disease, neuroinflammation, innate immunity, caspase-1, IL-1β, IL-18, pyroptosis

## Abstract

Neuroinflammation is a hallmark of several chronic neurodegenerative disorders, including Parkinson’s disease (PD) and other synucleinopathies, and is closely associated with microglial activation. The inflammasome adaptor protein apoptosis-associated speck-like protein containing a caspase activation and recruitment domain (ASC) undergoes oligomerization upon inflammasome activation, resulting in the formation of ASC specks. These structures are recognized by microglia as damage-associated molecular patterns (DAMPs), thereby promoting inflammatory signaling. Recent studies have demonstrated that extracellular ASC specks associate with amyloid-β (Aβ), and that uptake of ASC-Aβ composites by microglia amplifies pro-inflammatory responses. Although ASC specks have also been shown to potentiate α-synuclein (αSyn) preformed fibril-induced pathology, their interaction with monomeric αSyn and their effects on inflammasome activation remain poorly understood. Here, we co-stimulated murine SIM-A9 microglia with ASC-αSyn assemblies and evaluated their effects on NLRP3 inflammasome activation. Electron microscopy revealed a close association between ASC fibrils and αSyn monomers. Exposure of microglia to ASC-αSyn enhanced NLRP3 inflammasome activation, as evidenced by increased ASC speck formation, caspase-1 activation, and secretion of IL-1β and IL-18, when compared to ASC or αSyn alone. In addition, immunohistochemical analyses of *post mortem* brain tissue from patients with dementia with Lewy bodies demonstrated ASC expression in regions exhibiting αSyn pathology and showed ASC co-localization with αSyn within Lewy bodies. Collectively, these findings identify ASC-αSyn assemblies as potent amplifiers of NLRP3 inflammasome signaling and suggest a mechanism through which they may contribute to sustained neuroinflammation in synucleinopathies.

## Introduction

Synucleinopathies are a group of heterogeneous neurodegenerative disorders characterized by progressive cell loss and aggregates of α-synuclein (αSyn) in neural cells in the form of Lewy bodies and glial cytoplasmic inclusions [1]. In addition to αSyn-rich Lewy bodies and Lewy neurites, co-pathologies, most notably of amyloid-β (Aβ)-containing plaques and phosphorylated tau-positive tangles, are frequently observed in synucleinopathies [2–4]. The most common ones are Parkinson’s disease (PD), dementia with Lewy bodies (DLB), and multiple system atrophy (MSA). αSyn has been shown to play a role in regulating neurotransmitter release via its presynaptic functions and lipid metabolism in neural cells that shapes synaptic plasticity [5]. Among the mechanisms implicated in disease progression, neuroinflammation has emerged as a major contributor. Evidence suggests a bidirectional relationship between αSyn and neuroinflammation, whereby αSyn can trigger inflammatory responses and, in turn, its metabolism is influenced by inflammatory processes and microbial infections [6–8].

The nucleotide-binding oligomerization domain (NOD)-, leucine-rich repeat (LRR)-, and pyrin domain (PYD)-containing protein 3 (NLRP3) inflammasome is a multiprotein complex that plays a central role in the innate immune system’s response to pathogens and cellular damage [9]. The inflammasome complex consists of the NLRP3 protein, the adaptor protein apoptosis-associated speck-like protein containing a caspase activation and recruitment domain (ASC), and pro-caspase-1. NLRP3 assembles the inflammasome complex by recruiting ASC and activating pro-caspase-1 through proximity-induced autoproteolysis. Active caspase-1 processes the pro-inflammatory cytokines interleukin (IL)-1β and IL-18 and cleaves gasdermin D to trigger pyroptotic cell death. Upon activation, NLRP3 assembles into the inflammasome complex and promotes formation of ASC specks, dense supramolecular structures that are approximately 1 μm in diameter [10, 11]. Beyond their role in inflammasome assembly, ASC specks can serve as scaffolds for protein aggregation and have been reported to possess an intrinsic ability to recruit and co-aggregate cytosolic proteins through hydrophobic interactions [12]. For example, extracellular ASC specks are biologically active and can cross-seed the aggregation of Alzheimer’s disease-associated Aβ in neighbouring microglia [13]. Furthermore, the uptake of ASC-Aβ composites by microglia triggers NLRP3 inflammasome activation and pyroptotic cell death, thereby resulting in the amplification of chronic pro-inflammatory responses through a self-propagating feed-forward cycle [14].

Studies on human tissues and preclinical models of PD have established a link between αSyn aggregation, microglial NLRP3 inflammasome activation and neuronal cell death [15]. Patients with PD exhibit increased expression of NLRP3, ASC, and caspase-1 within microglia of the *Substantia nigra pars compacta* (SNpc), as well as elevated extracellular ASC and circulating levels of NLRP3, caspase-1, and IL-1β [15, 16]. More recently, ASC specks were shown to amplify NLRP3 inflammasome activation induced by αSyn pre-formed fibrils (PFFs), enhancing reactive microgliosis, exacerbating αSyn pathology, dopaminergic neurodegeneration, and motor deficits [17]. However, the contribution of ASC specks and their interaction with monomeric αSyn in propagating inflammasome activation and amplifying inflammatory responses remain unclear. In the present study, we generated ASC-αSyn assemblies and investigated their effects on NLRP3 inflammasome activation in a murine SIM-A9 microglial cell culture model. In parallel, we also examined ASC distribution in *post mortem* human brain tissue affected by synucleinopathy to explore the potential association of ASC with αSyn *in vivo* and its relevance to neuroinflammatory responses.

## Materials and methods

### Ethics statement

Human brain tissues were obtained at autopsy at The Ottawa Hospital in Ottawa, Ontario, Canada. Ethics approval was obtained from the Ottawa Health Science Network Research Ethics Board (#20120963-01H). Brain tissue was processed in the Department of Pathology at The Ottawa Hospital according to routine procedures for *post mortem* brain collection.

### Cell culture

Mouse SIM-A9 microglial cells (ATCC; CRL-3265) were cultured in Dulbecco’s modified Eagle’s medium/Ham’s F12 50/50 Mix (DMEM/F12) (Wisent; 319-075-CL) containing 10% heat-inactivated Qualified Fetal Bovine Serum (FBS) (Gibco; 12483-020) under standard conditions at 37°C and 5% CO_2_. Cells were passaged every two to three days and cells between passage 16 and 30 were used for all experiments.

### Preparation of recombinant ASC

Human full-length ASC protein was cloned into a pET28a-derived expression vector containing a C-terminal Avitag and 6× Histidine tag and expressed in E. coli BL21. For protein expression, bacterial cultures were induced with 10 μM isopropyl β-D-1-thiogalactopyranoside (IPTG) and incubated overnight at 28°C. Cells were lysed by sonication and centrifuged to separate the lysate into pellet (insoluble protein) and supernatant (soluble protein) fractions. ASC was expressed at high yield and was found almost exclusively in insoluble inclusion bodies. ASC protein from the insoluble inclusion body fraction was solubilized in 8 M urea and refolded using standard protocols [18]. Refolded ASC was subsequently purified by size-exclusion chromatography in 20 mM Tris/HCl, 0.5 M L-arginine (pH 11) buffer.

To induce fibril formation, the ASC-containing solution was first buffer-exchanged into ice-cold phosphate-buffered saline (PBS) (Wisent; 311-010-CL) using an Amicon® Ultra 0.5mL centrifugal filter (10 kDa MWCO) (Millipore; UFC501024). Buffer exchange was performed by centrifugation at 12,000 × g for 10 min at 4°C with three successive washes. Throughout the procedure, samples were kept cold and a minimum retentate volume of >100 μL was maintained. ASC was transferred to LoBind Tubes (Eppendorf; 022431081), and ice-cold PBS was added to achieve the desired protein concentration. The protein solution was subsequently incubated at 37°C for 1 h to induce fibril formation, as described previously [14]. ASC fibrils were stored at 4°C and used within 2-3 days.

### Preparation of αSyn and Aβ_1-42_

Recombinant, wild-type human αSyn protein monomers (Type 1) were purchased at a stock concentration of 5 mg/ml in PBS (pH 7.4) (StressMarq Biosciences Inc.; SPR-321) and stored at −80°C. For *in vitro* treatments, αSyn was used at working concentrations of 2.5 µM and 5 µM. During preparation of working stocks, αSyn monomers were maintained on ice or at 4°C and were subjected to no more than two freeze/thaw cycles.

Human Aβ protein (1-42), pre-treated with 1,1,1,3,3,3-Hexafluoro-2-propanol (HFIP)-(Bachem AG; 4090148), was dissolved in sterile Dulbecco’s PBS (Gibco; 14040141) to create a stock concentration of 0.45 mg/ml and stored at −80°C. HFIP is a volatile solvent that disrupts Aβ fibrils and promotes monomer formation by destabilizing β-sheet structures and disrupting hydrophobic interactions within aggregated amyloid preparations [19]. For *in vitro* treatments, Aβ was used at a working concentration of 5 µM. During preparation of working stocks, Aβ monomers were maintained on ice or at 4°C and were subjected to no more than two freeze/thaw cycles.

### Generation of ASC-αSyn assemblies

Fibrillar ASC (2.5 μM) and αSyn monomers (2.5 μM) were incubated together overnight at 37°C in DMEM/F12 media supplemented with 1% FBS and 1% penicillin-streptomycin (P-S) (Wisent; 450-201-EL) in LoBind tubes. Single-protein control preparations were incubated under the same conditions in medium containing 1% FBS and 1% P-S. ASC-Aβ composites (1.75 μM ASC and 5 μM Aβ) were included as a positive control, as previously described [14].

### Electron microscopy

Fibrillary ASC (1.75 μM) and αSyn monomers (5 μM) were pre-incubated overnight at 37°C in Tris-buffered saline (TBS) (ThermoScientific; J60877.K2), as described above in the “*Generation of ASC-αSyn assemblies*” section. Four to six carbon support film copper grids (300 mesh) (Ted Pella, Inc.; 01843-F) were placed on a coverslip and glow-discharged using the PELCO easiGlowTM Glow Discharge System (Ted Pella, Inc.) in automatic mode.

For negative staining electron microscopy (EM), 5 μL of protein sample was applied to a glow-discharged copper grid and incubated for 2 min at room temperature. Excess liquid was removed using Whatman filter paper, and the sample was negatively stained with 5 μL of 2% uranyl acetate for 30 sec. Finally, residual stain was removed with filter paper, and the grids were allowed to air-dry completely. Samples were imaged using a JEM-1400Plus transmission electron microscope (JEOL) operated at 120 kV. Images were processed using ImageJ software (National Institute of Health).

### In vitro treatments

For *in vitro* experiments, mouse SIM-A9 microglia were seeded in 24-well plates at a density of 5 × 10^4^ cells/well in 500 μL DMEM/F12 supplemented with 10% FBS and allowed to adhere overnight. Inflammasome signaling was primed by treatment with 100 ng/mL lipopolysaccharide (LPS; from *E. coli* [O55:B5]) (Sigma; L6529) for 3 h. Following priming, cells were washed once with serum-free DMEM/F12 and treated with either 2.5 μM ASC, 2.5 μM αSyn, or ASC-αSyn (containing 2.5 μM ASC and 2.5 μM αSyn) for 24 h. After treatment, supernatants and cell lysates were collected for further analyses. Unless otherwise specified, all treatments were performed in DMEM/F12 media supplemented with 1% FBS and 1% P-S in a final volume of 200 μL per well.

### Immunocytochemistry and confocal microscopy

SIM-A9 microglia were seeded at a density of 5 × 10^4^ cells/well in 500 μL DMEM/F12 media supplemented with 10% FBS in 24-well plates containing poly-D-Lysine (PDL)-coated coverslips (100 μg/mL; Sigma-Aldrich; A-003-E). To confirm the microglial origin of mouse SIM-A9 microglia, rabbit anti-Iba-1 (1:500; Fujifilm Wako; 019-19741) and rabbit anti-CD68 (1:100; Abcam; ab303565) antibodies were used. Mouse anti-MAP2 (1:250; Sigma-Aldrich; M4403) and rabbit anti-GFAP (1:250; Dako; Z0334) antibodies were included as neuronal and astrocytic markers, respectively. All subsequent immunocytochemical procedures were performed as described below.

Treatments were performed as described previously. Following treatments, cells were washed once with PBS and fixed in 4% paraformaldehyde (Biotium; 22023) prepared in PBS for 15 min at room temperature. Cells were then washed once with PBS and permeabilized with 0.25% Triton X-100 for 10 min. Cells were blocked with Dako Protein Block, Serum-Free (Agilent Technologies, Inc.; X0909) for 20 min. Primary antibodies were diluted in Dako Antibody Diluent (Agilent Technologies, Inc.; S0809) and incubated overnight at 4°C in a humidified chamber. To assess NLRP3 activation and ASC speck formation, mouse anti-NLRP3/NALP3 (1:100; Cryo-2; AdipoGen; AG-20B-0014-C100) and rabbit anti-ASC (1:100; clone AL177; AdipoGen; AG-25B-0006-C100) antibodies were used, respectively. After three washes with PBS, cells were incubated for 30 min with either Alexa Fluor™ 488 goat anti-mouse (1:500; Invitrogen; A-11029) or Alexa Fluor™ 488 goat anti-rabbit (1:500; Invitrogen; A-11008). After three additional PBS washes, Phalloidin-iFluor 594 Reagent (Abcam; ab176757) was applied at a dilution of 1:2,000 together with the goat anti-rabbit secondary antibody to visualize F-actin filaments. Coverslips were quickly rinsed once in dH_2_O, blotted perpendicularly on Kimwipes and mounted onto microscope slides using Dako Fluorescent Mounting Medium (Agilent Technologies, Inc.; S3023) spiked with 5 μg/mL Hoechst 33258. To assess caspase-1 activity, the FAM-FLICA® Caspase-1 (YVAD) Assay Kit (ImmunoChemistry Technologies LLC; 98) was used according to the manufacturer’s protocol. Cells were mounted on microscope slides, allowed to air dry for a few minutes, and imaged immediately.

All images were captured using a Leica STELLARIS 5 Confocal Microscope (Leica Microsystems) with Z-stack imaging (z-step of 0.8 μm) using either a 20× or 63× objective. Every image was imaged from the farthest point to the closest point from the objective. Image processing was performed using LAS X Office (Leica Microsystems) and image analysis was conducted using ImageJ (National Institute of Health). Quantitative analyses were performed on 150-1200 cells per biological replicate using 20× images only. Total cell counts and NLRP3 mean fluorescence intensity were determined using automated image analysis, whereas ASC specks and active caspase-1-positive cells were quantified manually.

### Immunohistochemistry and immunofluorescence of human post mortem brain tissues

Following tissue collection at autopsy, brain specimens were fixed in 20% neutral buffered formalin for a minimum of 10-14 days. The midbrain was dissected, processed, and embedded in paraffin. 5 µm thick sections were cut, dried overnight, deparaffinized in xylene (Fisher Chemicals; X54), and rehydrated through a graded series of decreasing ethanol concentrations.

For chromogenic immunohistochemistry, staining was performed using a Bond-Max fully automated system (Leica Biosystems, Wetzlar, Germany). ASC immunoreactivity was detected using a modified protocol J and the Bond Polymer Red Refine Detection Kit (Leica Biosystems; DS9390), followed by detection of αSyn using a modified protocol F and the Bond Polymer Refine Detection Kit (Leica Biosystems; DS9800). Following deparaffinization and rehydration, sections underwent heat-induced epitope retrieval in Epitope Retrieval Solution 1 (citrate buffer, pH 6; Leica Biosystems; AR9961) at 98°C for 10 min. Endogenous peroxidase activity was quenched with peroxidase block for 5 min. Sections were then incubated with anti-ASC antibody (1:500; Santa Cruz, sc-514414) for 20 min at room temperature. After washing, sections were sequentially incubated with post-primary alkaline phosphate (AP) reagent for 20 min and polymer AP for 20 min, followed by mixed Red Refine detection in a two-step process (10 min and 5 min). Following ASC detection, sections were incubated with anti-αSyn antibody (1:10,000; StressMarq; SMC-600) for 10 min at room temperature. After washing, sections were incubated with post-primary reagent for 8 min and polymer reagent for 8 min, followed by mixed DAB (3, 3’-diaminobenzidine) Refine solution for 10 min. Sections were counterstained with hematoxylin for 6 min, dehydrated, cleared, and coverslipped using Micromount mounting media (Leica Biosystems). Slides were air-dried overnight prior to imaging. Brightfield mages were acquired using an Olympus IX81 microscope equipped with a colour camera.

For immunofluorescence staining, endogenous peroxidase activity was quenched with 3% hydrogen peroxide prepared in PBS for 15 min. Heat-induced antigen retrieval was then performed with citrate buffer (pH 6) for 20 min. To reduce non-specific binding, sections were incubated in Dako Serum-Free Protein Block (Agilent Dako; X0909) for 1 h at room temperature. Double immunofluorescence labeling was performed sequentially. Following removal of excess blocking solution, sections were incubated overnight at 4°C with anti-ASC antibody (1:100; Santa Cruz; sc-514414). After washing, sections were incubated with biotinylated goat anti-mouse secondary antibody (1:250; Vector Laboratories; BA-9200) for 45 min at room temperature, followed by streptavidin-CY3 (1:1000; Jackson Immunoresearch; 106-160-084) for 30 min to visualize ASC immunoreactivity.

Sections were subsequently incubated overnight at 4°C with one of the following primary antibodies: anti-αSyn (1:500; StressMarq; SMC-600), anti-Iba-1 (1:500; Synaptic Systems; 234009), or anti-tyrosine hydroxylase (TH; 1:200; Pel-Freeze; P40101). Following washing, appropriate secondary antibodies were applied for 45 min at room temperature. Alexa Fluor™ 488 goat anti-rabbit IgG (1:500; Invitrogen; A-11034) was used to detect α-Syn and TH, while Alexa Fluor™ 488 goat anti-chicken IgG (1:500; Invitrogen; A-11039) was used to detect Iba-1. All primary and secondary antibodies were diluted in Dako antibody diluent (Agilent Dako).

Slides were coverslipped using ProLong Glass Antifade Mountant with NucBlue (Invitrogen) and allowed to dry overnight in the dark. Fluorescent images were acquired using a Leica Stellaris 5 laser-scanning confocal microscope (Leica Biosystems, Wetzlar, Germany). Single optical sections were captured using LAS X software (version 3.10.0.28982; Leica Microsystems), with identical laser power, detector gain, and pinhole settings maintained across all experimental conditions.

### Western blotting

Protein concentrations of cell lysates were determined using the DC Protein Assay (Bio-Rad; 5000112) according to the manufacturer’s instructions.

To confirm the presence of recombinant human ASC and αSyn and to assess ASC oligomer formation, protein preparations were treated with 5 mM disuccinimidyl suberate (DSS) at 37°C for 30 min to stabilize ASC oligomers through irreversible cross-linking. Samples were then analyzed by SDS-PAGE and Western blotting using anti-ASC (1:1,000; AdipoGen; AG-25B-0006-C100) and anti-αSyn (1:1,000; Abcam; ab27766) antibodies.

For cellular analyses, SIM-A9 microglia were seeded at 2.2 × 10^5^ cells/well in 6-well plates and treated as described in the “*in vitro* treatments” section. Following treatment, cells were collected and lysed in 1× radioimmunoprecipitation assay (RIPA) buffer (Sigma; R0278) supplemented with protease and phosphatase inhibitors (Thermo Fisher Scientific; 1861280). Lysates were denatured in 4× Laemmli sample buffer (Bio-Rad; 1610747) for 5 min at 95°C, and 20–40 μg of protein was separated on 4–20% Mini-PROTEAN TGX™ precast gels (Bio-Rad; 4561093) and transferred to PVDF membranes (Bio-Rad; 1620177). Membranes were stained with Ponceau S (ThermoScientific; A40000279) to verify protein transfer and normalize protein loading, blocked in 5% non-fat dry milk, and incubated overnight at 4°C with primary antibodies against NLRP3 (1:1,000; AdipoGen; AG-20B-0014-C100), ASC (1:1,000; AdipoGen; AG-25B-0006-C100), caspase-1 (1:1,000; AdipoGen; AG-20B-0042), and GSDMD (1:1,000; Abcam; ab209845). Following incubation with HRP-conjugated secondary antibodies for 1 h at room temperature, immunoreactive bands were detected using SuperSignal™ West Pico PLUS Chemiluminescent Substrate (Thermo Fisher Scientific; 34577) and imaged with a ChemiDoc™ Imaging System (Bio-Rad). Densitometric analysis was performed using Image Lab software (Bio-Rad). Protein expression was normalized to total lane protein determined by Ponceau S staining and expressed relative to untreated controls as fold change.

### Enzyme-linked immunosorbent assay (ELISA)

Following treatment of SIM-A9 microglia, cell culture supernatants were collected and analyzed for IL-1β and IL-18 levels using commercially available mouse IL-1β (R&D Systems; DY401-05) and IL-18 (R&D Systems; DY7625-05) ELISA kits, according to the manufacturer’s protocols using Immulon 4 HBX Flat Bottom Plates (ThermoScientific; 3855). Colour development was achieved using TMB High Sensitivity ELISA Substrate (Abcam; ab171523) for 7 min and stopped with 2 N H_2_SO_4_. Absorbance was measured at 450 nm with background correction at 540 nm using a microplate reader.

### Cell death assay

Cell death was quantified by measuring lactate dehydrogenase (LDH) release using the Cytotoxicity Detection KitPLUS (LDH) (Roche; 04744926001). SIM-A9 microglia were treated as described above in the “*in vitro treatments*” section, and LDH levels were measured according to the manufacturer’s protocol.

### Statistical analyses

Data are presented as mean ± standard error of mean (SEM). Statistical significance was determined using an ordinary one-way analysis of variance (ANOVA) followed by Tukey’s multiple comparisons test where the means of the test columns was compared individually to the mean of every other column. Normal (Gaussian) distribution was assumed for all analyses. ASC-Aβ composites (containing 1.75 μM ASC) were not statistically compared with ASC alone or ASC-αSyn (containing 2.5 μM ASC) because the treatments contained different amounts of oligomerized ASC. Asterisks denote statistical significance levels as follows: p < 0.05 (*), p < 0.01 (**), p < 0.001 (***), and p < 0.0001 (****). All statistical analyses were performed using GraphPad Prism 9.0 (GraphPad Software Inc., San Diego, CA, USA).

## Results

### Monomeric αSyn clusters around ASC fibrils

ASC specks possess an intrinsic ability to promote protein aggregation on their surface [12], and extracellular, microglia-derived ASC specks have been demonstrated to associate and bind to Aβ monomers [13]. To determine whether ASC specks may similarly associate with αSyn, we first produced recombinant human ASC protein by large scale expression of ASC in *E. coli* BL21 cells. Western bot analysis demonstrated that ASC was present predominantly in its monomeric form with an apparent molecular weight of approximately 25 kDa (**Figure 1A**).

**Figure 1.**
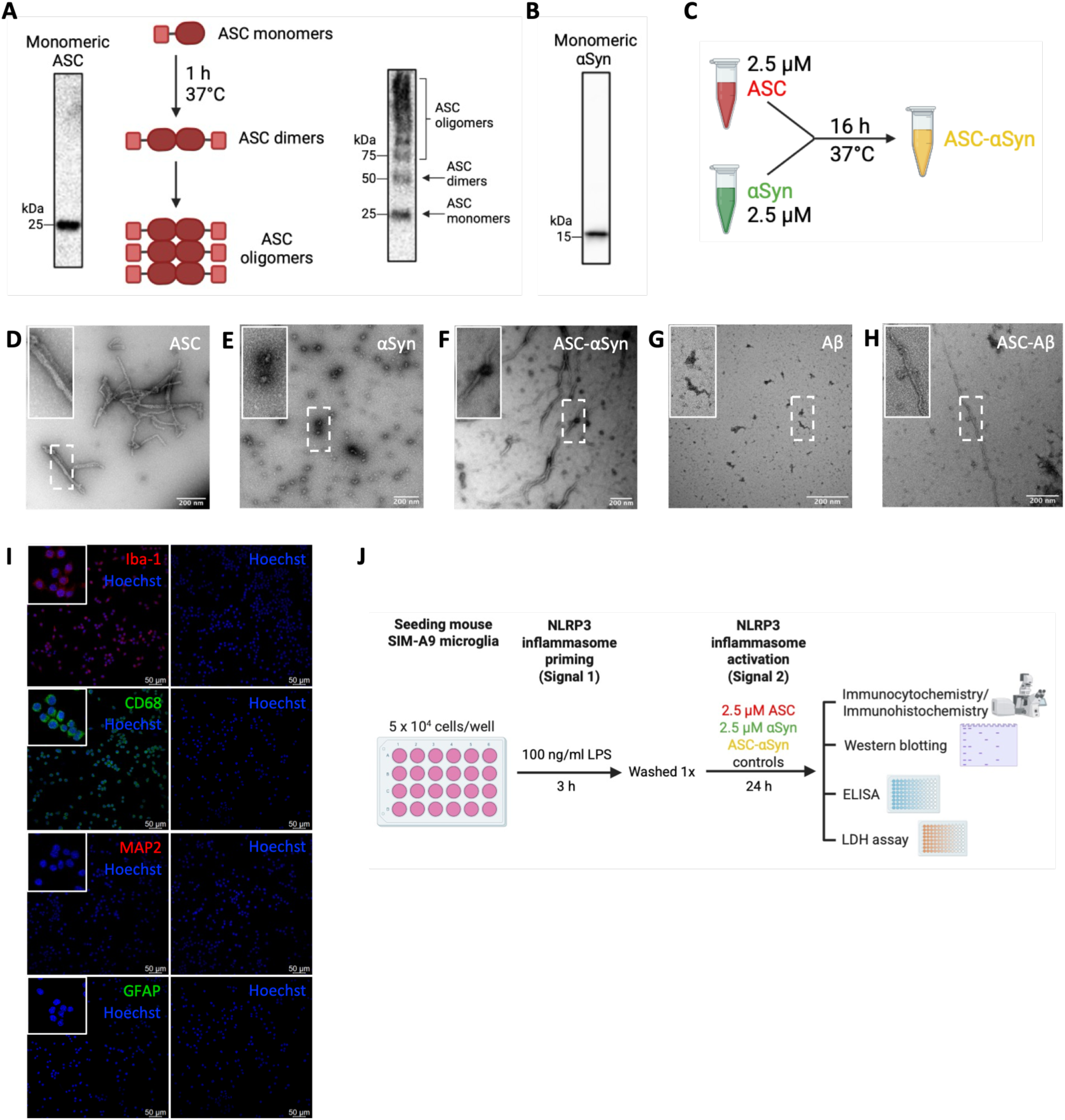
Generation of ASC-αSyn assemblies and characterization of the SIM-A9 microglial model. (A) Schematic illustration of ASC oligomerization. Recombinant human ASC was incubated at 37°C for 1 h to induce oligomer formation and analyzed by SDS-PAGE and Western blotting. Immunoblotting with an anti-ASC antibody identified ASC monomers (∼25 kDa), dimers (∼50 kDa), and higher-order oligomers (>50 kDa). (B) Western blot analysis confirming the presence of recombinant human monomeric αSyn. (C) Schematic illustration of ASC-αSyn assembly generation. Oligomerized ASC (2.5 µM) and monomeric αSyn (2.5 µM) were co-incubated at 37°C for 16 h. (D-H) Transmission electron microscopy images of ASC (D), αSyn (E), ASC-αSyn (F), Aβ (G) and ASC-Aβ (H). ASC-Aβ composites served as a positive control. Scale bars, 200 nm. (I) Immunocytochemical characterization of SIM-A9 microglia. Cells were positive for Iba-1 and CD68, microglia/macrophage markers, and negative for the neuronal marker, MAP2, and astrocytic marker, GFAP. The right panel shows secondary antibody-only controls. Scale bars, 50 µm. (J) Schematic overview of the experimental design used for *in vitro* inflammasome activation studies. Figure panels A-C and J were created with BioRender.com.

To induce oligomerization, purified ASC protein monomers were incubated at 37°C for 1 h to promote ASC ‘prionoid’ activity, as previously described [14]. Following incubation, Western blot analysis confirmed the presence of ASC monomers (∼25 kDa), dimers (∼50 kDa), and higher-order oligomers (>50 kDa) (**Figure 1A**). Oligomerized ASC was subsequently co-incubated with human monomeric αSyn (**Figure 1B**) at a 1:1 molar ratio for 16 h at 37°C to form ASC-αSyn assemblies (**Figure 1C**). Cells treated with ASC-Aβ composites were included as a positive control, as previously described [14].

To characterize the resulting protein assemblies, samples were analyzed by negative stain electron microscopy (EM). ASC was seen to arrange into long, slightly twisted helical filaments with a relatively uniform surface morphology (**Figure 1D**). In contrast, αSyn alone appeared largely monomeric and lacked detectable fibrillar structures (**Figure 1E**). In co-incubated samples, αSyn monomers were found in close proximity to ASC fibrils (**Figure 1F**). As expected, Aβ monomers (**Figure 1G**) were observed to be in association with ASC fibrils, forming ASC-Aβ composites (**Figure 1H**).

### ASC-αSyn co-stimulation activates the NLRP3 inflammasome in SIM-A9 microglia

To investigate the effects of ASC-αSyn treatment on NLRP3 inflammasome activation, we first confirmed the suitability of mouse SIM-A9 cells as a microglial cell model. SIM-A9 cells, a spontaneously immortalized murine microglial cell line [20], expressed the microglia/macrophage markers ionized calcium-binding adapter molecule 1 (Iba-1) and cluster of differentiation 68 (CD68), but lacked immunoreactivity for the neuronal marker microtubule-associated protein 2 (MAP2) and the astrocytic marker glial fibrillary acidic protein (GFAP) (**Figure 1I**) [21].

To assess NLRP3 inflammasome activation, SIM-A9 microglia were primed with 100 ng/ml LPS for 3 h and stimulated with 2.5 μM ASC, 2.5 μM αSyn, or ASC-αSyn (2.5 μM ASC: 2.5 μM αSyn) for 24 h (**Figure 1J**). As expected, immunofluorescence analyses demonstrated that LPS treatment alone increased NLRP3 protein expression compared with untreated controls, consistent with inflammasome priming (**Figure 2A-B**). Following LPS priming, treatment with ASC, αSyn, or ASC-αSyn further increased NLRP3 protein expression (**Figure 2A-B**).

**Figure 2.**
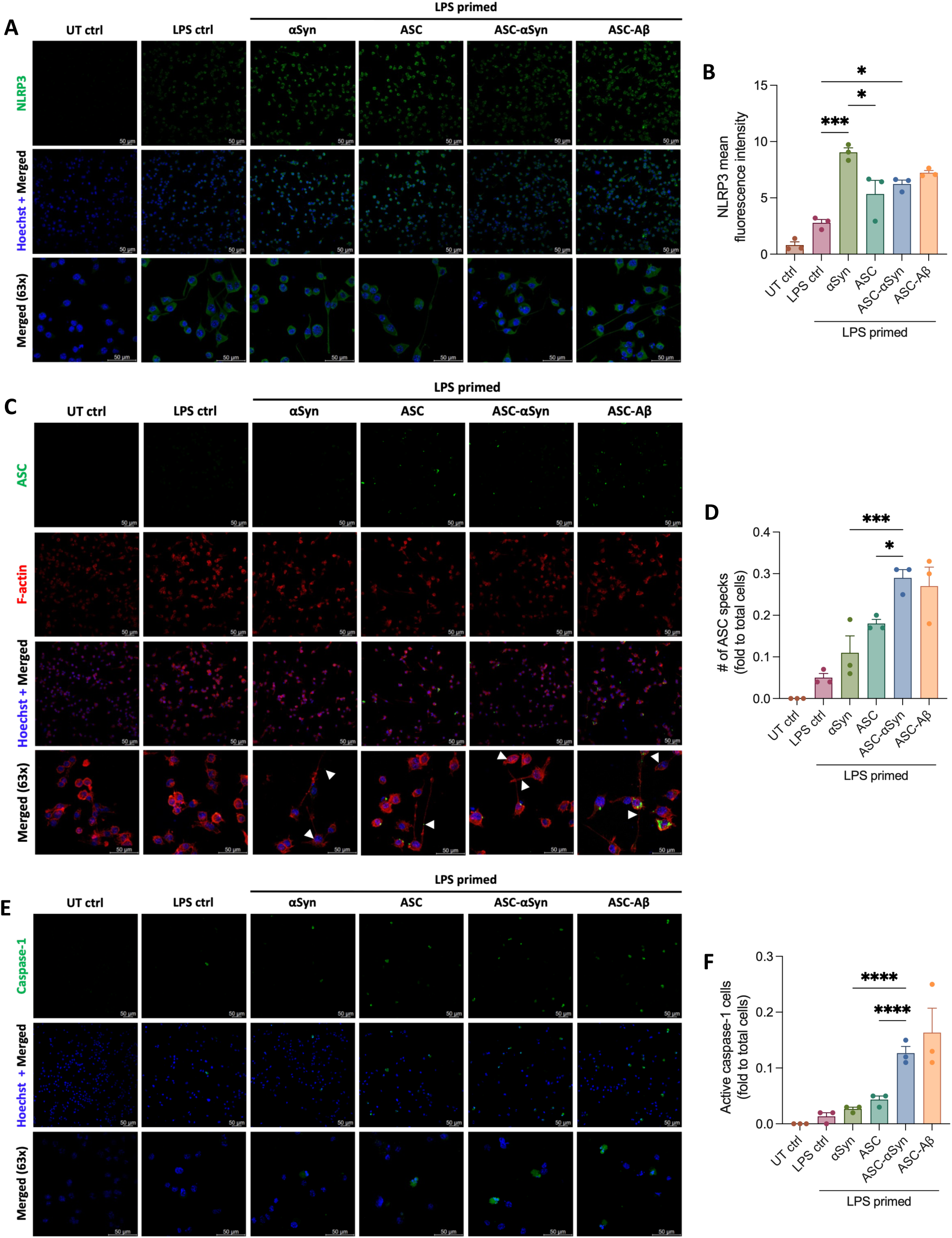
ASC-αSyn treatment enhances NLRP3 inflammasome activation in SIM-A9 microglia. Cells were primed with 100 ng/ml LPS for 3 h and subsequently treated with ASC, αSyn, or ASC-αSyn for 24 h. Immunocytochemical detection and quantification of NLRP3 protein (A–B), ASC speck formation (C-D), and active caspase-1-positive cells (E-F). Arrowheads indicate endogenous ASC specks (∼1 µm in diameter). Images were acquired using 20× and 63× objectives. Scale bars, 50 µm. Data is representative of three independent experiments (n = 3) and presented as mean ± SEM. Statistical analyses were performed using one-way ANOVA followed by Tukey’s post-hoc multiple comparisons test. Significance levels are indicated as follows: *p < 0.05, ***p < 0.001, ****p < 0.0001. Asterisks indicate significance between groups.

Upon inflammasome activation, ASC binds to NLRP3 to create the inflammasome complex and interacts with other ASC proteins to assemble into an ASC speck. This ASC speck formation has been used as a readout for inflammasome activation [22]. Compared with the LPS control, ASC-αSyn significantly increased the number of ASC specks (*p* < 0.0001; **Figure 2C-D**). Moreover, ASC-αSyn-treated cells exhibited more ASC specks than cells treated with ASC (*p* = 0.0273) or αSyn (*p* = 0.0009) alone, respectively (**Figure 2C-D**).

To determine whether ASC speck formation was accompanied by downstream inflammasome signaling, caspase-1 activity was assessed after 24 h of ASC-αSyn treatment. Immunofluorescence analyses revealed increased caspase-1 activity in ASC-αSyn treated cells compared with ASC (*p* < 0.0001) and αSyn alone (*p* < 0.0001; **Figure 2E-F**). ASC-αSyn-treated cells also displayed greater fluorescent intensity, suggesting enhanced caspase-1 activation (**Figure 2E**). Notably, activated caspase-1-positive cells exhibited morphological features consistent with pyroptosis, including nuclear condensation, membrane ballooning, and membrane rupture (**Figure 2E**) [23]. Together, these findings indicate that ASC-αSyn stimulation of microglia promote NLRP3 inflammasome activation and downstream caspase-1 induction.

To further evaluate the effects of ASC-αSyn treatment on NLRP3 inflammasome activation, protein levels of NLRP3, full-length caspase-1 and GSDMD were analyzed by Western blotting (**Figure 3A**). Consistent with the immunofluorescence data, LPS priming increased NLRP3 protein levels in comparison to untreated controls (**Figure 3B**). In parallel to an increase in active caspase-1-positive cells (**Figure 2I**), a reduction in the full-length caspase-1 protein was observed, suggesting caspase-1 processing and activation (**Figure 3C**). ASC-αSyn-treated cells exhibited a significant, 1.3-fold decrease in full-length caspase-1 compared to LPS-treated controls (*p* = 0.0442), whereas ASC-treated cells showed a similar trend that did not reach significance (*p* = 0.0921; **Figure 3C**). Increased levels of cleaved GSDMD were detected following ASC and ASC-αSyn stimulation compared with LPS- and αSyn-treated cells (no statistical significance; **Figure 3D**), consistent with enhanced pore-forming activity downstream of caspase-1 activation.

**Figure 3.**
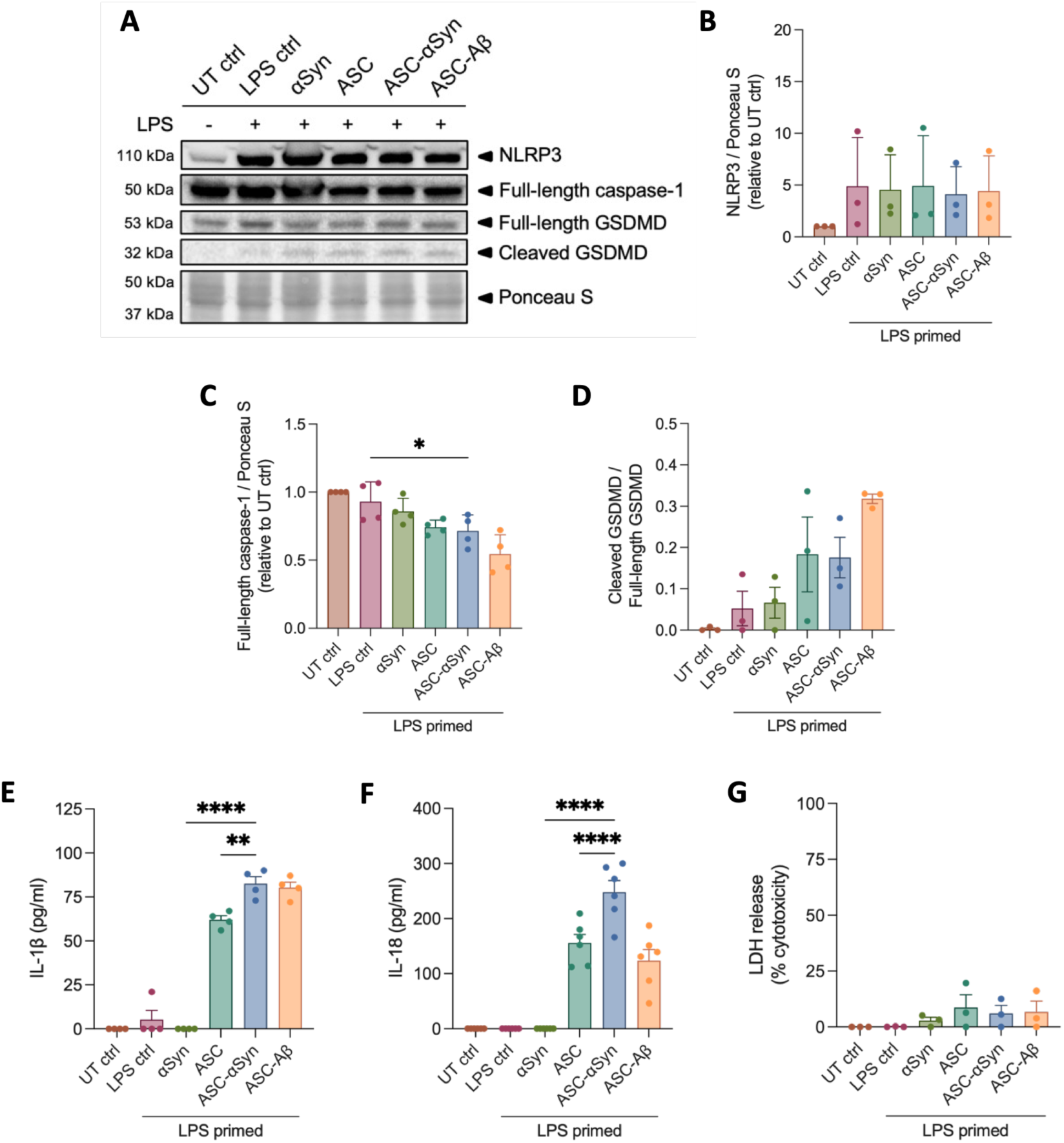
ASC-αSyn treatment amplifies NLRP3 inflammasome signaling and proinflammatory cytokine release in SIM-A9 cells. (A) Representative Western blots of NLRP3, full-length caspase-1, and full-length and cleaved gasdermin D (GSDMD) in SIM-A9 microglia primed with 100 ng/mL LPS for 3 h and subsequently treated with ASC, αSyn, or ASC-αSyn for 24 h. (B–E) Densitometric quantification of NLRP3 (B), full-length caspase-1 (C), full-length GSDMD (D), and cleaved GSDMD (E). Protein levels were normalized to total protein staining (Ponceau S) and expressed as fold change relative to untreated (UT) controls. (F–H) Quantification of IL-1β (F), IL-18 (G), and lactate dehydrogenase (LDH; H) levels in conditioned media collected from SIM-A9 microglia following LPS priming and treatment with ASC, αSyn, or ASC–αSyn for 24 h. Data were obtained from four independent experiments for NLRP3, caspase-1, and IL-1β analyses (B, C, F; n = 4), six independent experiments for IL-18 analysis (G; n = 6), and three independent experiments for GSDMD and LDH analyses (D, E, H; n = 3). Data are presented as mean ± SEM. Statistical analyses were performed using one-way ANOVA followed by Tukey’s multiple-comparisons test. Significance levels are indicated as follows: *p < 0.05, **p < 0.01, ****p < 0.0001. Asterisks indicate significance between groups.

IL-1β and IL-18 secretion in cell culture supernatants were quantified by ELISA as indicators of caspase-1 activity [24]. ASC-αSyn induced a significant increase in IL-1β release compared with αSyn alone (*p* < 0.0001) or ASC alone (*p* = 0.0025; **Figure 3E**). Similarly, ASC-αSyn significantly increased IL-18 release compared with ASC alone (*p* < 0.0001; **Figure 3F**). In contrast, treatment with LPS or αSyn alone did not induce detectable amounts of IL-1β or IL-18 release under the experimental conditions. These findings suggest that ASC-αSyn assemblies enhance pro-inflammatory cytokine release beyond that observed following exposure to ASC or αSyn alone.

To determine whether ASC-αSyn-induced inflammasome activation affected cell viability, LDH release was measured as an indicator of cell death. Despite LPS priming and ASC-αSyn stimulation, LDH release remained low across all treatment groups, suggesting minimal cell death during the treatment period (**Figure 3G**). Together, these findings suggested to us that ASC-αSyn activate the NLRP3 inflammasome, caspase-1 signaling, and pro-inflammatory cytokine release in microglia to a greater extent than when compared to ASC and αSyn alone.

### ASC co-localizes with αSyn pathology in human midbrain tissue

Previous studies have reported up-regulated ASC expression and of ASC specks formation in the SNpc of PD patients [15]. In addition, Lewy bodies isolated from *post mortem* PD brains have been shown to positively stain for both caspase-1 and αSyn [25]. To determine whether ASC is similarly associated with αSyn pathology, we examined ASC and αSyn (pSer129) expression in sections of *post mortem* human midbrain. Compared with healthy controls, DLB cases exhibited elevated ASC immunoreactivity, consistent with increased inflammasome-associated signaling and inflammation (**Figure 4**). In contrast, healthy control tissues displayed minimal ASC and αSyn immunoreactivity (**Figure 4A, C**). Immunohistochemical and immunofluorescence analyses further revealed that Lewy bodies were positively stained for both ASC and αSyn (pSer129), demonstrating their co-localization within these pathological inclusions (**Figure 4B, D**).

**Figure 4.**
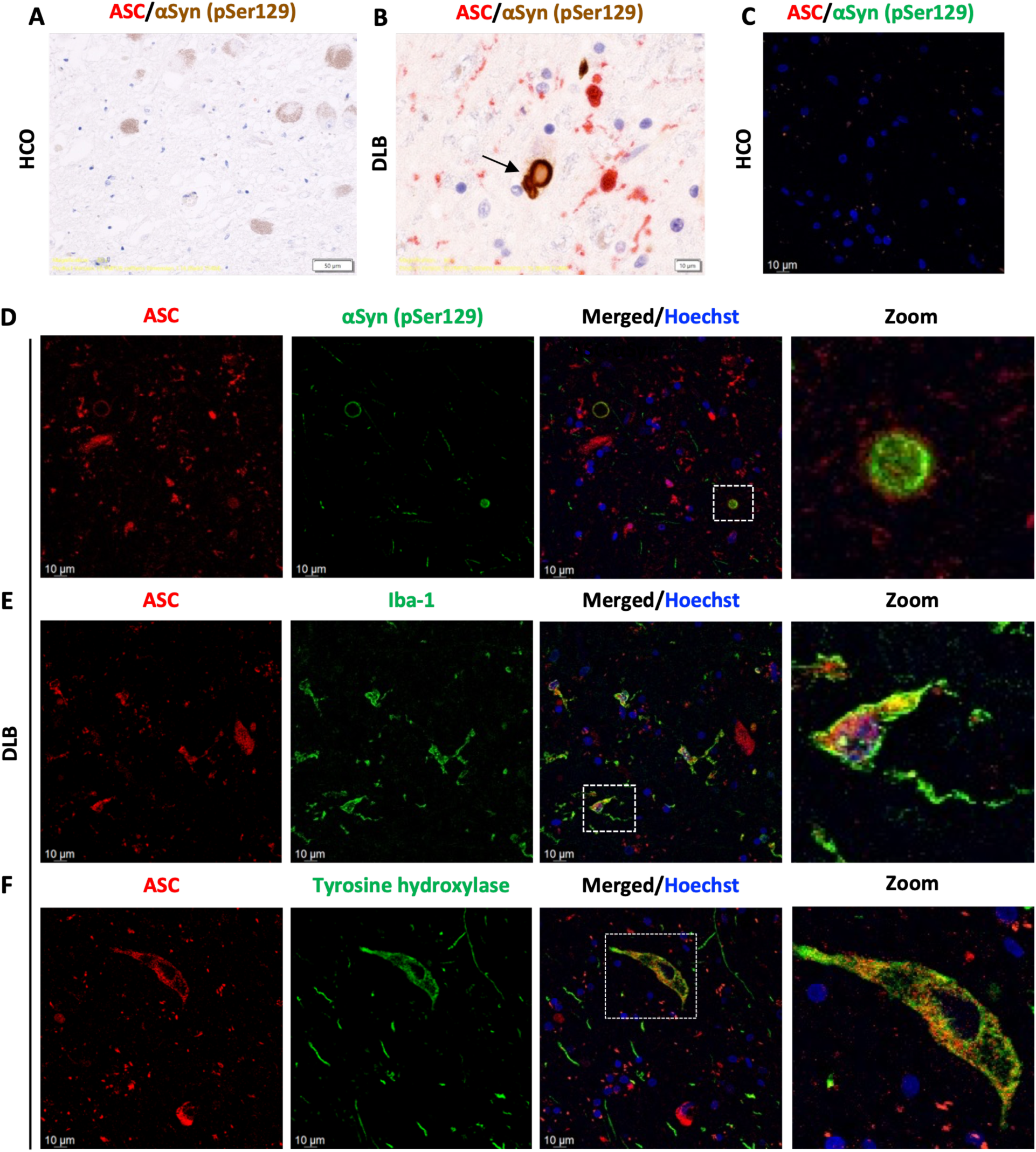
ASC co-localizes with αSyn pathology in the human midbrain. (A, B) Immunohistochemical staining of ASC and αSyn in *post mortem* midbrain tissue from healthy controls (HCO, n = 2) and patients with dementia and Lewy bodies (DLB, n = 5). (C-D) Immunofluorescence analysis of αSyn and ASC in HCO and DLB tissue. (E-F) Immunofluorescence analyses demonstrating co-localization of ASC with Iba-1-positive microglia (E) and co-localization of ASC with tyrosine hydroxylase (TH)-positive dopaminergic neurons (F) in DLB tissue. In panel B, the arrow indicates a Lewy body positive for both ASC and αSyn. In panel D, boxed region is shown at higher magnification to highlight co-localization of ASC and αSyn within a Lewy body. In panels E-F, boxed regions are shown at higher magnification to illustrate ASC co-localization with Iba-1, and TH-positive dopaminergic neurons, respectively. Scale bars, 10 μm.

In addition to its localization within Lewy bodies, ASC was detected in myeloid cells and dopaminergic neurons in diseased brain tissue exhibiting αSyn pathology. In DLB brain tissue, ASC co-localized with Iba-1, consistent with previous reports demonstrating ASC expression in myeloid cells during neuroinflammation and inflammasome activation [26, 27] (**Figure 4E**). Because αSyn preferentially accumulates in dopaminergic neurons and contributes to their degeneration, we next investigated whether ASC was also associated with tyrosine hydroxylase (TH)-positive neurons. TH is the rate-limiting enzyme in catecholamine synthesis and a well-established marker of dopaminergic neurons [28, 29]. Consistent with a recent report demonstrating elevated ASC expression in dopaminergic neurons in PD [27], ASC co-localized with TH-positive dopaminergic neurons in diseased *post mortem* brain tissue (**Figure 4F**). Collectively, these findings demonstrate that ASC is associated with αSyn pathology and localizes to both Lewy bodies and TH-positive dopaminergic neurons in the human midbrain. These observations further support a link between ASC and αSyn-related neuropathology.

## Discussion

Neuroinflammation, characterized by the presence of excessive levels of inflammatory markers in neural tissues, is increasingly recognized as a key contributor to the pathogenesis of PD and related synucleinopathies [30, 31]. Among the inflammatory pathways, implicated in disease progression, the NLRP3 inflammasome has emerged as a central mediator of innate immune responses through activation of caspase-1, maturation of the IL-1 family cytokines and induction of pyroptotic cell death. A growing body of evidence implicates αSyn aggregation, microglial NLRP3 inflammasome activation, and extracellular ASC specks in the pathogenesis of PD and related synucleinopathies [15, 32]. However, whether ASC specks directly influence αSyn-associated inflammatory responses remain incompletely understood. In the present study, we demonstrate that oligomerized ASC associates with exogenous monomeric αSyn, potentiates NLRP3 inflammasome activation *in vitro* and augments inflammatory responses in microglia. Furthermore, we show increased ASC levels and co-localization of ASC and αSyn in human midbrain tissue exhibiting Lewy body pathology.

ASC is an adaptor protein that bridges the NLRP3 sensor and pro-caspase-1 effector during inflammasome assembly. Upon activation, ASC assembles into large supramolecular structures known as ASC specks, which serve as signaling platforms that promote caspase-1 activation and inflammatory cytokine maturation [22, 33, 34]. Beyond their intracellular role, ASC specks exhibit prion-like properties and can persist extracellularly following pyroptosis [32, 35, 36]. Extracellular ASC specks have been shown to retain biological activity, recruit pro-caspase-1, and propagate inflammatory responses in neighbouring cells [32, 37–39]. Moreover, microglia-derived ASC specks have been reported to bind Aβ and accelerate amyloid deposition in Alzheimer’s disease models [13]. Consistent with these observations, we reproduced the association between ASC and Aβ in our positive control experiments and extended these findings to αSyn (**Figure 5**). EM revealed that monomeric αSyn clustered along ASC fibrils following co-incubation, suggesting a physical association between the two proteins (**Figure 1D**). Although the structural basis of this interaction remains unknown, these observations raise the possibility that ASC fibrils may serve as scaffolds capable of recruiting aggregation-prone proteins. Such a mechanism would be consistent with the ability of ASC specks to promote Aβ aggregation [13] and could represent a broader role for ASC in proteinopathies characterized by protein accumulation (**Figure 5**). Although our findings do not establish a direct molecular binding between ASC and αSyn, they demonstrate a close spatial association that warrants further investigation using complementary biophysical approaches.

**Figure 5.**
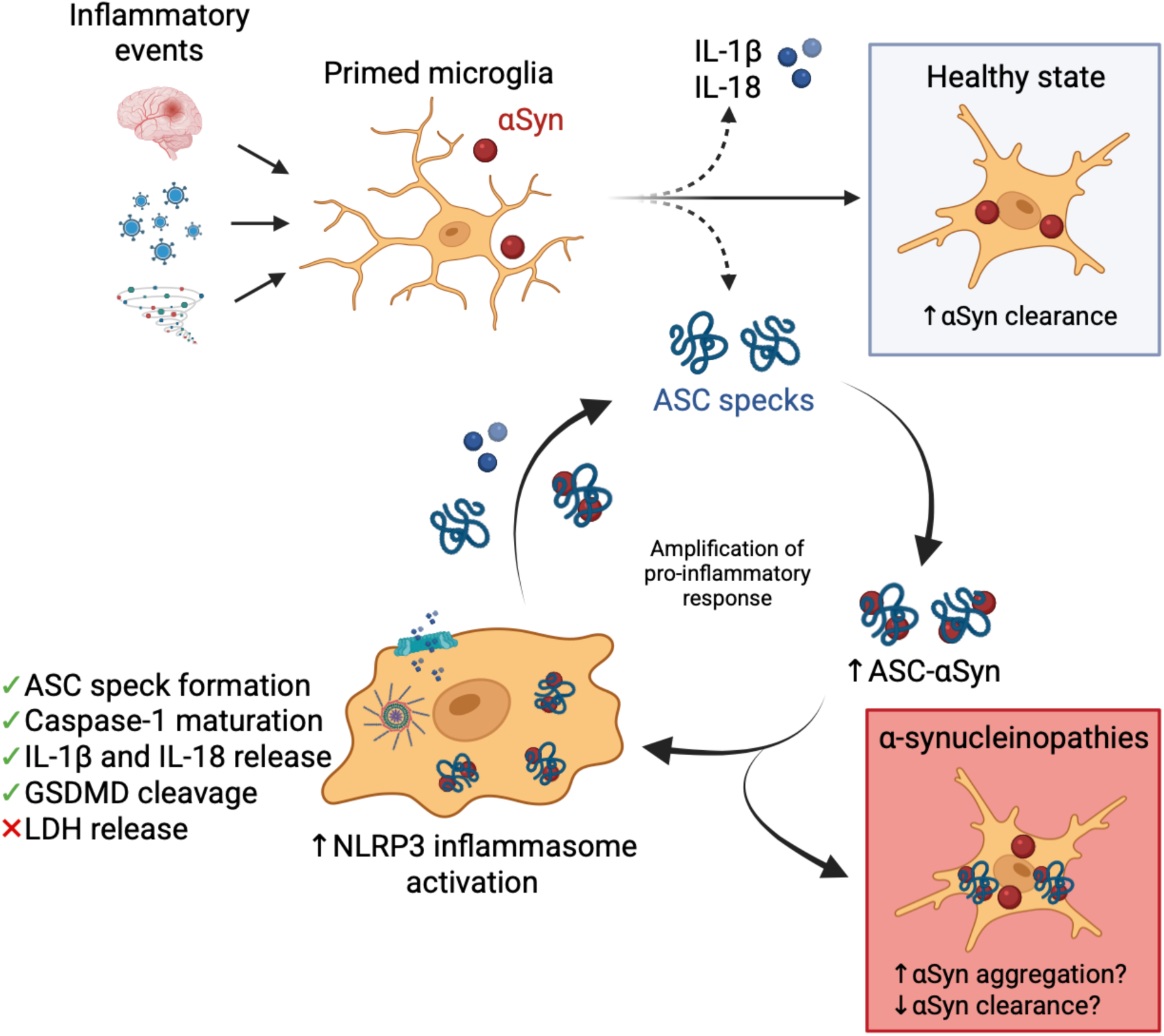
Proposed model illustrating the contribution of ASC-αSyn to amplification of microglial inflammatory responses. Different conformations of αSyn and inflammatory stimuli promote microglial priming and activation through pattern recognition receptors and other cell surface receptors. Under physiological conditions, activated microglia internalize extracellular αSyn and contribute to its clearance. Following NLRP3 inflammasome activation, inflammasome-associated components, including ASC specks and the mature pro-inflammatory cytokines IL-1β and IL-18, are released into the extracellular space. Extracellular ASC specks may associate with αSyn and form ASC-αSyn assemblies. Uptake of ASC-αSyn by microglia may further enhance NLRP3 inflammasome activation, leading to increased ASC speck formation, caspase-1 activation, and cytokine release. Extracellular ASC specks may also recruit soluble ASC and interact with extracellular αSyn, thereby contributing to the propagation of inflammatory signaling. Collectively, these events may establish a feed-forward cycle that amplifies neuroinflammation and contributes to the progression of αSyn pathology. Figure created with BioRender.com.

To investigate the biological consequences of this association, we examined inflammasome activation in LPS-primed microglia. Age-associated increases in basal inflammatory signaling, often referred to as ‘inflammageing’ is believed to lower the threshold for NLRP3 inflammasome activation and may contribute to the increased susceptibility to neurodegenerative diseases observed in older individuals [40–42]. Following priming, ASC-αSyn induced greater inflammasome activation than ASC or αSyn alone. This was evidenced by increased ASC speck formation, elevated caspase-1 activity, enhanced GSDMD cleavage, and increased secretion of IL-1β and IL-18 (**Figures 2 and 3**). Despite evidence of inflammasome activation, LDH release remained low across treatment groups. These findings indicate that ASC-αSyn assemblies primarily promoted inflammatory signaling under the experimental conditions tested and that inflammasome activation was not accompanied by substantial cytotoxicity *per se* within the 24-hour treatment period. Similar dissociation between inflammasome activation and overt cell death have been reported in microglia and other myeloid cell populations, where inflammatory cytokine release can occur in the absence of extensive membrane rupture [43–45]. Together, these findings suggest that the presence of αSyn in association with ASC amplifies inflammatory signaling downstream of NLRP3 activation. Notably, ASC specks were observed both intracellularly and extracellularly (**Figure 2C**). Extracellular ASC assemblies retain their biological activity and can be internalized by neighbouring myeloid cells [32], thus their persistence may facilitate the propagation of inflammatory responses beyond the initially activated cell population. In the context of synucleinopathies, extracellular ASC specks may encounter αSyn released from stressed or damaged neurons, thereby creating conditions favoring propagation of neuroinflammation.

The relationship between extracellular αSyn and innate immune activation has received considerable attention in recent years. Although αSyn is predominantly a cytosolic neuronal protein, both monomeric and oligomeric forms can be released into the extracellular environment through unconventional secretion pathways and in response to neuronal activity or injury [46–50]. Further, monomeric αSyn has been reported to induce inflammasome priming-associated responses without supplying a sufficient, second activation signal [51, 52]. Once extracellular, αSyn can be taken up by microglia and activate inflammatory signaling pathways [47–49]. Previous studies have largely focused on aggregated or pathogenic forms of αSyn as triggers of inflammasome activation. Our findings suggest that even non-fibrillar αSyn may contribute to inflammatory responses when associated with ASC assemblies. This observation may be particularly relevant during early disease stages, when extracellular αSyn species may be recruited into heteromeric assemblies and aggregates, leading to reduced levels of its monomers in human cerebrospinal fluid and interstitial fluid [53–56], as intracellular Lewy body pathology develops.

*Post mortem* brain tissues from PD patients and preclinical PD animal models have revealed increased ASC levels and the formation of ASC specks in the SNpc of PD patients and striatum of mice [15, 57, 58]. In biomarker studies, gene expression levels of ASC have been found to be elevated in peripheral blood mononuclear cells of PD patients compared to healthy control individuals [58]. In contrast, C-terminal truncation of αSyn can increase exposure of the non-amyloid-β component (NAC) region, thus promoting a pro-aggregatory conformational state [59]. Truncation of the C-terminal has been estimated to occur in 10-30% of total αSyn within Lewy bodies [60]. Caspase-1 may truncate αSyn in neurons through direct cleavage at Asp121 to induce αSyn aggregation and inflammation, further aggravating cytotoxicity and creating a positive-feedback inflammatory loop [61–63]. In addition, Lewy bodies isolated from *post mortem* PD brains were positively stained for both caspase-1 and αSyn, substantiating their co-existence in Lewy bodies and supporting the belief that caspase-1 may be involved in generating the truncated αSyn observed in Lewy bodies [25]. Based on the size of the Lewy body core and what is known about the ASC speck’s ability to cross-seed protein aggregation, it has been speculated that ASC specks can likely be found in the core of Lewy bodies comparably to Aβ plaques [13, 64]. Here, we corroborated increased ASC levels in cases of αSyn pathology and further revealed co-localization of ASC to Lewy bodies (**Figure 4**). Although these findings do not establish a causal role for ASC in Lewy body formation, they support the notion that inflammasome-derived ASC may contribute to the pathological microenvironment in which αSyn aggregation can occur or in which such aggregates are sequestered (**Figure 5**).

Our study has several limitations. First, while EM demonstrated close spatial association between ASC fibrils and αSyn monomers, the biophysical determinants of this interaction remain unknown. Future studies will be required to determine whether αSyn directly binds ASC and to identify the domains involved. Second, not all αSyn or Aβ monomers appeared associated with ASC fibrils, suggesting that only a subset of protein molecules participate in assembly formation under the conditions tested. Finally, using exogenous recombinant ASC may influence the visualization and quantification of endogenous ASC specks in microglial cells. To minimize this, endogenous ASC specks were quantified manually based on their characteristic size and intracellular localization.

In conclusion, our findings identify a previously underappreciated association between ASC specks and αSyn that enhances microglial inflammasome activation. The observation that ASC associates with αSyn *in vitro* and co-localizes with Lewy body pathology in human brain tissue further supports a link between inflammasome signaling and synucleinopathies. Our findings raise the possibility that extracellular ASC assemblies contribute to the propagation of neuroinflammation during disease progression in PD and DLB and position ASC as a potential molecular link between innate immune activation and neuronal synucleinopathies. Future studies investigating the molecular mechanisms underlying ASC-αSyn interactions and their impact on neuronal integrity may provide important insights into the early pathological mechanisms and related targets to potentially modify the course of neurodegeneration.

## Author Contributions

Conceptualization: K. T., C. P., M. G. S., J. K. S. Data acquisition: K.T., C.P., M.H., K.B., and D.C. Analysis and Writing – original draft: K.T. Writing – review & editing: K. T., C. P., D.C., J. M. W., J. J. T., M. G. S., J. K. S. Funding acquisition: C. P., M. G. S., J. K. S.

## Acknowledgments

We would like to thank Dr. Jyh-Yeuan (Eric) Lee (University of Ottawa) to use his laboratory for EM sample preparation, and The Transmission Electron Microscopy Core (University of Ottawa) for their valuable assistance with EM imaging. M.G.S. is supported by the Bhargava Family Research Chair in Neurodegeneration at The Ottawa Hospital.

